# From village to globe: A dynamic real-time map of African fields through PlantVillage

**DOI:** 10.1101/2019.12.19.882936

**Authors:** Annalyse Kehs, Peter McCloskey, John Chelal, Derek Morr, Stellah Amakove, Bismark Plimo, John Mayieka, Gladys Ntango, Kelvin Nyongesa, Lawrence Pamba, Melodine Jeptoo, James Mugo, Mercyline Tsuma, Winnie Onyango, David Hughes

**Affiliations:** Department of Entomology and Biology, Pennsylvania State University, University Park, PA, United States; Department of Agriculture and Biotechnology, Moi University, Eldoret, Kenya; Plant Production and Protection Division, United Nations Food and Agriculture Organization, Rome, Italy; Office of the Associate CIO for Research, Pennsylvania State University, University Park, PA, United States

**Keywords:** Earth Observations, Sentinel, Ground-truthed, polyculture, crowdsourcing

## Abstract

A major bottleneck to the application of machine learning tools to satellite data of African farms is the lack of high-quality ground truth data. Here we describe a high throughput method using youth in Kenya that results in a cost-effective method for high-quality data in near real-time. This data is presented to the global community, as a public good, on the day it is collected and is linked to other data sources that will inform our understanding of crop stress, particularly in the context of climate change.

## Introduction

A major goal of the global community interested in leveraging Earth Observations for smallholder farmers is to identify the composition of crops in farmers’ fields. A near real-time database of the location and type of crops that are grown would be very helpful for multiple stakeholders. These range from Governments and markets wishing to predict yield to epidemiologists and Earth scientists interested in crop-specific predictions of pests and climate change effects at the field level (McNally-2019). The importance of Earth Observations (EO) in achieving the United Nations Sustainable Development Goals was recently emphasized (Whitcraft et al-2019).

Researchers have successfully leveraged Earth Observations to identify crop composition in the United States (US) and the European Union (EU) farms (McNally-2019). In those settings, fields tend to be monocropped and the boundaries are regular with uniform planting dates throughout a region. This is not the case for smallholder farms in African countries. Intercropping and an abundance of weeds are very common. In addition, trees are often present inside the fields which proves a challenge for accurate satellite detection of crops. Further, the field boundaries are highly irregular and the low availability of seed and necessary farm inputs means fields might be only partially sown or sown with several different varieties.

While the remote observation of African farms with satellites would appear challenging with respect to farms elsewhere in the world, the need is certainly great (Whitcraft et al-2019). Hunger is on the rise in almost all subregions of Africa, where the prevalence of undernourishment has reached levels of 22.8 percent in sub-Saharan Africa (FAO-2019). Invasive pests like the Fall Armyworm (Day-2017) and severe changes in weather (Giordano-2019) associated with climate change are making a bad situation worse (Global Report on Food Crises-2019).

It is possible that high-resolution satellite data such as the 4-meter resolution Dove satellites^1^ from Planet or the 0.46-meter resolution satellites from Maxar would help identify the crops within fields^2^. However, their high cost and low flyover rate preclude their utility for a cross-continental dynamic map which is updated each season as crops change. Therefore, the pragmatic approach likely requires the use of free, open-access satellite data such as the European Space Agency’s (ESA) Copernicus system or the US National Aeronautics and Space Administration’s (NASA) LandSat data.

If the medium resolution satellites like Sentinel and LandSat are to be effectively used in the African smallholder farms context then it is very clear that accurate ground truth data, vital to validating predictive models, is the rate-limiting step. Moreover, the ground truth data must be open access to allow the global community to identify where the successes and shortcomings lie. Here we describe a high throughput method of accurate ground truth data collection from a smallholder farm setting in Kenya that not only results in high-quality data but the availability of the data for the global community on the same day it was collected through PlantVillage^3^. Further, we demonstrate how our data can be combined with other relevant data streams such as the United Nations Food and Agricultural Organization’s (UN FAO) Water Productivity through open-access of remotely sensed derived data Tool^4^ (WaPOR) and the African Soil Health Initiative. Our approach has the added value of directly employing young African scientists and building capacity in African science.

## Methods

### Mapping agents

We conducted in-field surveys in Busia County, Kenya throughout May 2019. We will refer to this time period throughout the rest of the paper as season one due to our collection being in the middle of Kenya’s first growing season of the calendar year (Place-2006). Busia County is one of the 47 counties of Kenya. The county has a land area of 1,694.5 (km^2^) and has an estimated population of 743,946 persons from the 2009 census (Busia County Integrated Development Plan-2018). “The County is situated at the extreme Western region of Kenya and borders Bungoma to the North, Kakamega to the East and Siaya to the South East, Lake Victoria to the South West and the Republic of Uganda to the West. It lies between latitude 0º and 0º 45 North and longitude 34º 25 East (Busia County Integrated Development Plan-2018)”. We developed a digital survey application using the Open Data Kit^5^ Build site. Field surveys were first performed by a team of two researchers who travelled from the USA to Busia, Kenya.

In August 2019 we implemented an alternative method which engaged Kenyan youth. We will refer to this time period throughout the rest of the paper as season two due to the data collection being in Kenya’s second growing season of the calendar year (Place-2006). We targeted youth because of the recognized high unemployment in Kenya and other African countries (FAO-2018). We were interested in determining whether collecting ground truth data and conducting in-field surveys could be a viable job option for Kenyan youth. We identified and hired 10 recent graduates in Agriculture from Moi University in Eldoret, Kenya (authors #4-14 of this paper). Members of the Kenyan survey team were paired together based on three main criteria; gender, background/experience and personality, with respective order of importance. Each pair comprised of one male and female. Both persons within the pair studied in a different program at Moi University (Agricultural Extension Education or Agricultural Biotechnology) to encourage collaboration between the two and complement the strengths/weaknesses of the other. One week of training was conducted to educate the 10 graduates on how to use the ODK Collect application to map fields and conduct in-field surveys. The pairs stayed locally within the communities of the regions they were assigned. Each member of the Kenyan survey team was paid a daily allowance of $14/day for salary, $4/day for transport, $2.5/week for call data, and $20/month for 15GB of internet data. The salary includes accommodation costs and is within the rates of internship allowance offered by local companies and institutions. The transport allowance covers the cost to move with each pair on one motorbike with appropriate safety equipment each day. Once arrived at their destination, field to field movements were done on foot. The call data was used for communication to the lead farmers, following farmers and seed entrepreneurs. The mobile data connection was used to upload images and surveys to a shared Google Drive account and for daily communication with the team in the United States via WhatsApp.

We had pre-existing connections in the community through an Irish charity called Self Help Africa (SHA) which had a Lead Farmer and Following Farmer model with each Lead Farmer having between 20 - 35 Following Farmers. Based on this existing network, the Kenyan survey team were distributed evenly across Busia county such that each pair of the Kenyan survey team was based near a different cluster of SHA Lead Farmers. The Kenyan survey team worked with the Lead Farmers in their region, who would then introduce them to additional community members with fields.

### ODK Application and Data Collection Protocol

The ODK form is built using ODK Build^5^ and is used to survey fields in conjunction with the ODK Collect application. The app is available to others upon request. The user of the form collects the following information from the farmer for each field to be mapped: GPS location of the field, field boundaries, crop type and the ratio of different crops, the time and date of survey, planting date, harvest date, density of crop, disease prevalence for cassava mosaic virus, brown streak virus, and green mites, and maize fall armyworm, five pictures of the field including at least one landscape overview picture of the field, and any comments. Comments can describe any relevant information with regards to the field. For example, if the field had to be replanted due to lack of rain or low germination rates, or if they used any control methods for diseases or pests.

When the survey pair arrive at the field, the protocol is to open ODK Collect application and begin to fill out a blank survey form. To fill out a blank survey, the farmer is required to be present to answer questions such as when the crop was planted, when it will be harvested, crop variety, if the field was fertilized, etc. Next, a GPS point and shape of the field are collected. There are three options to map the shape of the field. The first input method, which was used in May, is ‘Placement by tapping’. This method requires a fair/good cellular data connection to connect and download satellite imagery sourced from Google. This method also assumes a familiarity with interpreting satellite maps. The second method, ‘Manual location recording’, requires the user to walk to each vertex of the field and capture a GPS coordinate manually. The third method, ‘Automatic location recording’, requires that the user walks around the boundary of the field while the application automatically collects location coordinates. The location coordinates are recorded every 20 seconds with an accuracy of ±3 meters. Methods two and three do not require a data connection or any previous experience in mapping fields. The Kenyan survey team began mapping the shape of fields using methods two and three until they became experienced at reading satellite images, which occurred within two months of beginning their work. Method three is continually used when the data connection is not strong or their data connection is roaming due to proximity to Uganda. After collecting the shape of the field, the point location information and the farmer has answered the rest of the survey questions, the form is finalized and saved.

### Local Storage of ODK Survey Forms

During season one, the finalized survey forms were submitted by pulling the forms manually off the phone with ODK Briefcase and aggregated into a comma-separated values file (CSV). The CSV’s were then uploaded to Google Drive to store and subsequently shared with others. We also stored the CSV on an external hard drive as a backup. In season two, we developed a new method for form submission. This method also uses Google Drive as the submission platform and for storage, however, the finalized forms are uploaded directly to a shared Google Sheet using a data connection. The images captured within the ODK are stored in Google Drive as well. Google Drive has limits on the amount of data that can be stored within Google Sheets. As such, it is a temporary step. The long-term storage of the data is in the PlantVillage database, which is an S3 Bucket in AWS. The global community can access this data via the PlantVillage website^3^. The data and associated metadata can be downloaded as a CSV file upon request.

### Privacy

We ask farmers for permission to collect the data from their farms and permission to use that data to improve our understanding of negative effects like pests and climate change. When we visit farms the only privileged data is the name of the farmer and whether she owns the land. This data is not shared. We did not encounter obstacles to sharing data. The farmers we engaged with clearly understood the twin threats of pests and climate change and wanted ways to tackle them. We provide farmers with advice both when we collected data on their fields and later on through an SMS alert system to repay them for sharing the data.

### Water Productivity via WaPOR

The UN FAO has developed a database called the Water Productivities Open-access portal^4^. This tool provides a series of measurements on water use by crops for Africa and the Middle East (FAO and IHE Delft-2019). We retrieve the WaPOR data for the locations which have been mapped using the open source WaPOR API. This provides added value to the community as the data on water productivity are presented alongside the crop composition data to give a crop stress measurement. The crop stress is determined by comparing the actual to reference evapotranspiration.

### The Dynamic PV Map

The surveyed fields, disease diagnosis data captured from PlantVillage *Nuru*, and WaPOR data can all be found on the PlantVillage website^3^. This page combines the surveyed fields with disease diagnosis information captured in the PlantVillage app to determine hotspots of disease. The WaPOR data is added to the map using the filter. When the WaPOR filter is on, it shows a green, amber, or red circle over a disease diagnosis marker that represents the crop stress.

### Ethics Statement

This paper does not contain any studies involving animals performed by any of the authors. This paper does not contain any studies involving human participants performed by any of the authors. Ethical approval was not required for the recruitment of the ten graduate students as they are scientific collaborators(authors #4-14 on this paper).

## Results

### Collection rate

In season one, 474 fields were surveyed over the course of the month (6^th^ May to 3^rd^ June). The field collection rate was 26 fields/day. The median fields collected per day was 26.5 with a mode of 32 (Figure 1).

**Figure 1:**
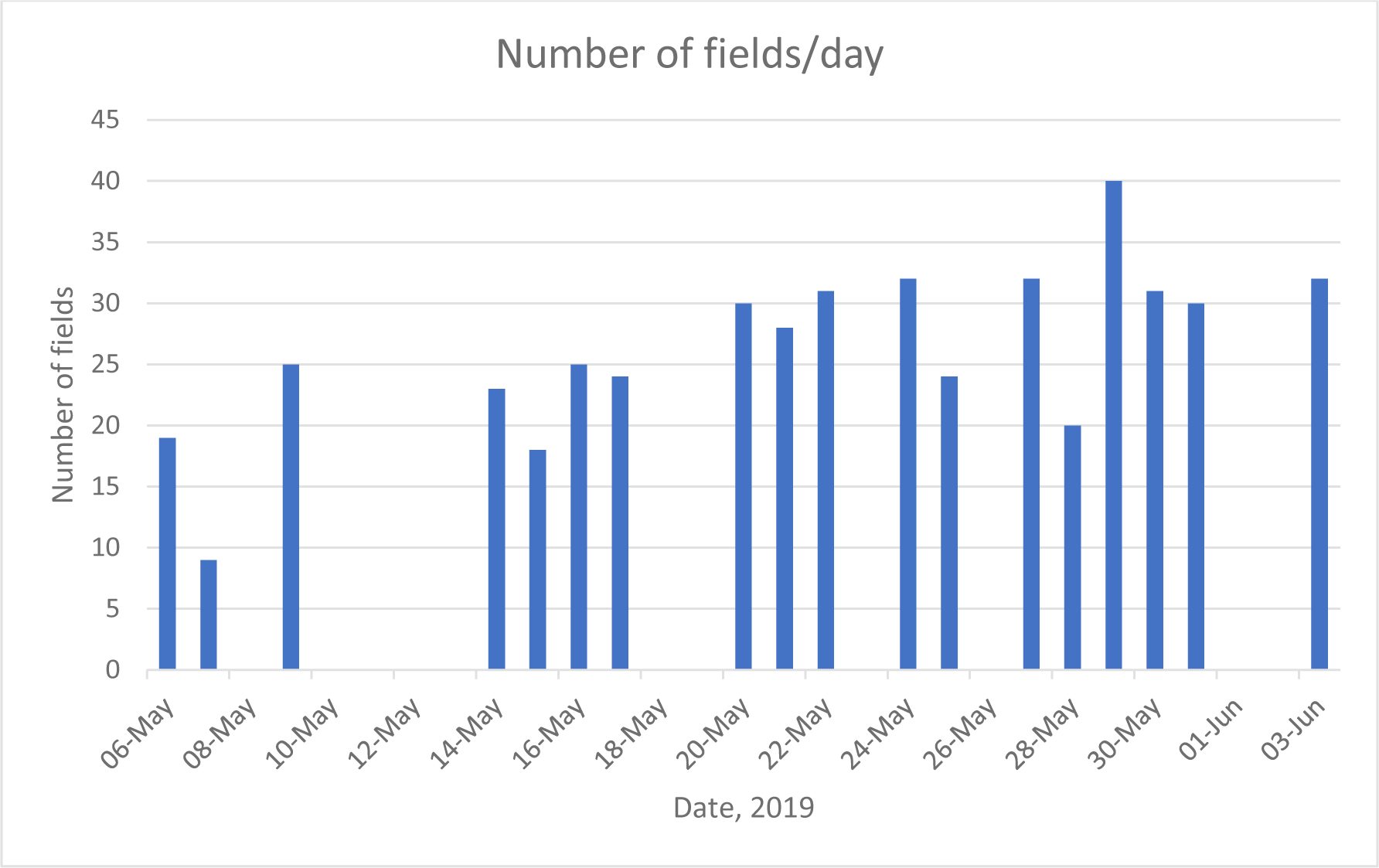
Number of fields surveyed each day between May 06^th^ and June 03^rd^, 2019 in Busia, Kenya

In season two we began field collections with the Kenyan survey team. The total number of fields collected between August 13^th^ and November 21^st^ was 9,263. The collection rate was 128 fields/day which equates to 25.6 fields per pair. The median fields collected per day is 137.5 with a mode of 143 fields per day (Figure 2).

**Figure 2:**
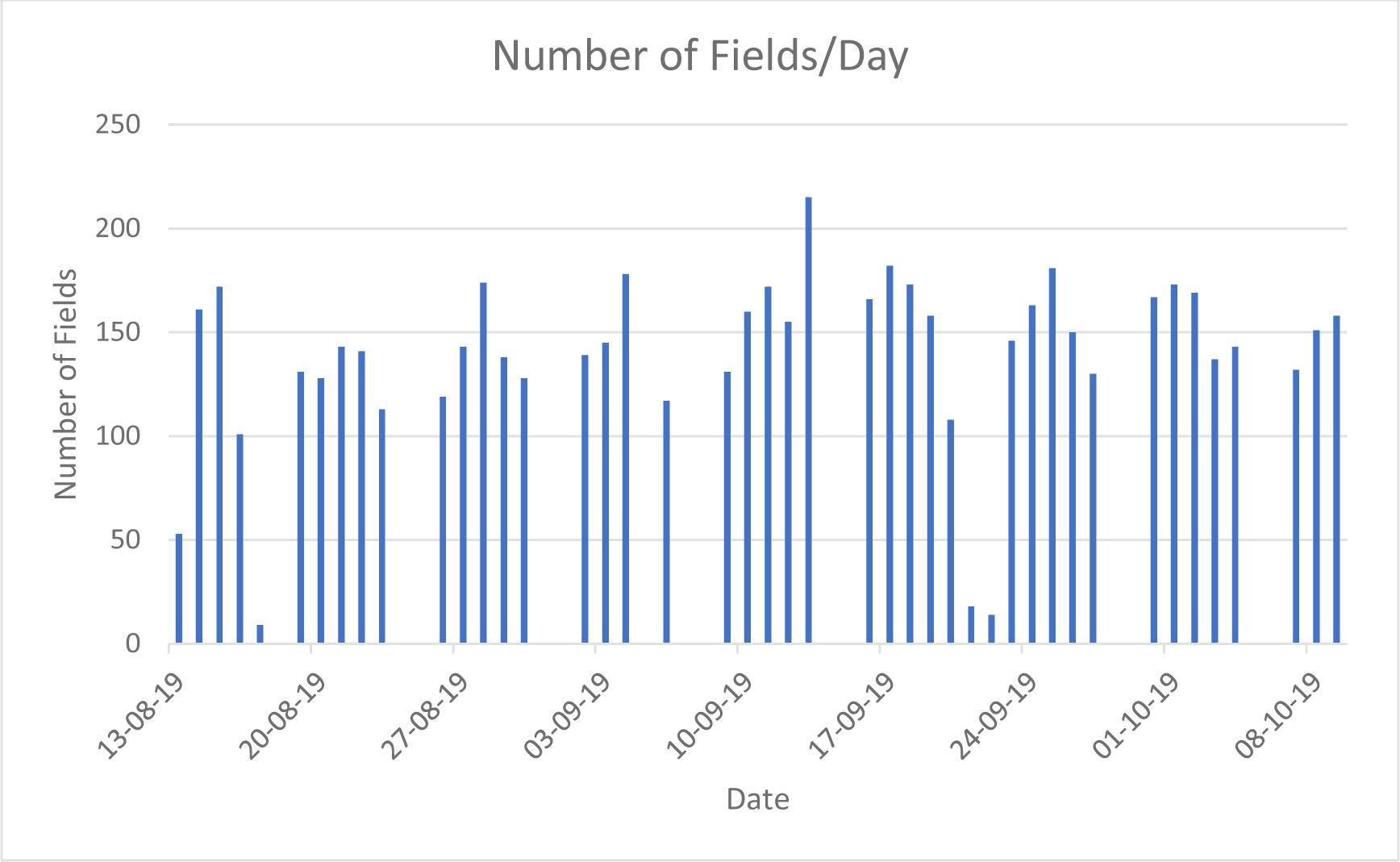
Number of fields surveyed each day between August 13^th^ and November 21^st^ in Busia, Kenya

### Composition of Fields

For both the season one and season two collections, our surveys showed that fields were a combination of intercrop and monocrop settings. Intercropped fields were divided into two categories: intercropped with one other crop and intercropped with greater than one other crop. The breakdown of the fields collected from August 13^th^ to November 21^st^, 2019 by the Kenyan survey team is 65% monocrop and 35% intercrop. Of the intercropped fields collected, 85% are intercropped with one other crop and 15% are intercropped with greater than one other crop (Figure 3). The crop category labeled ‘Not Listed’ encompasses any crops that were not included in the 10 crops that were determined to be the majority of crops planted in Kenya. Some of the crops that fall into the ‘Not Listed’ category include sweet potato, cotton, cowpea, sesame, kale, tomato, Napier grass, banana and others.

**Figure 3:**
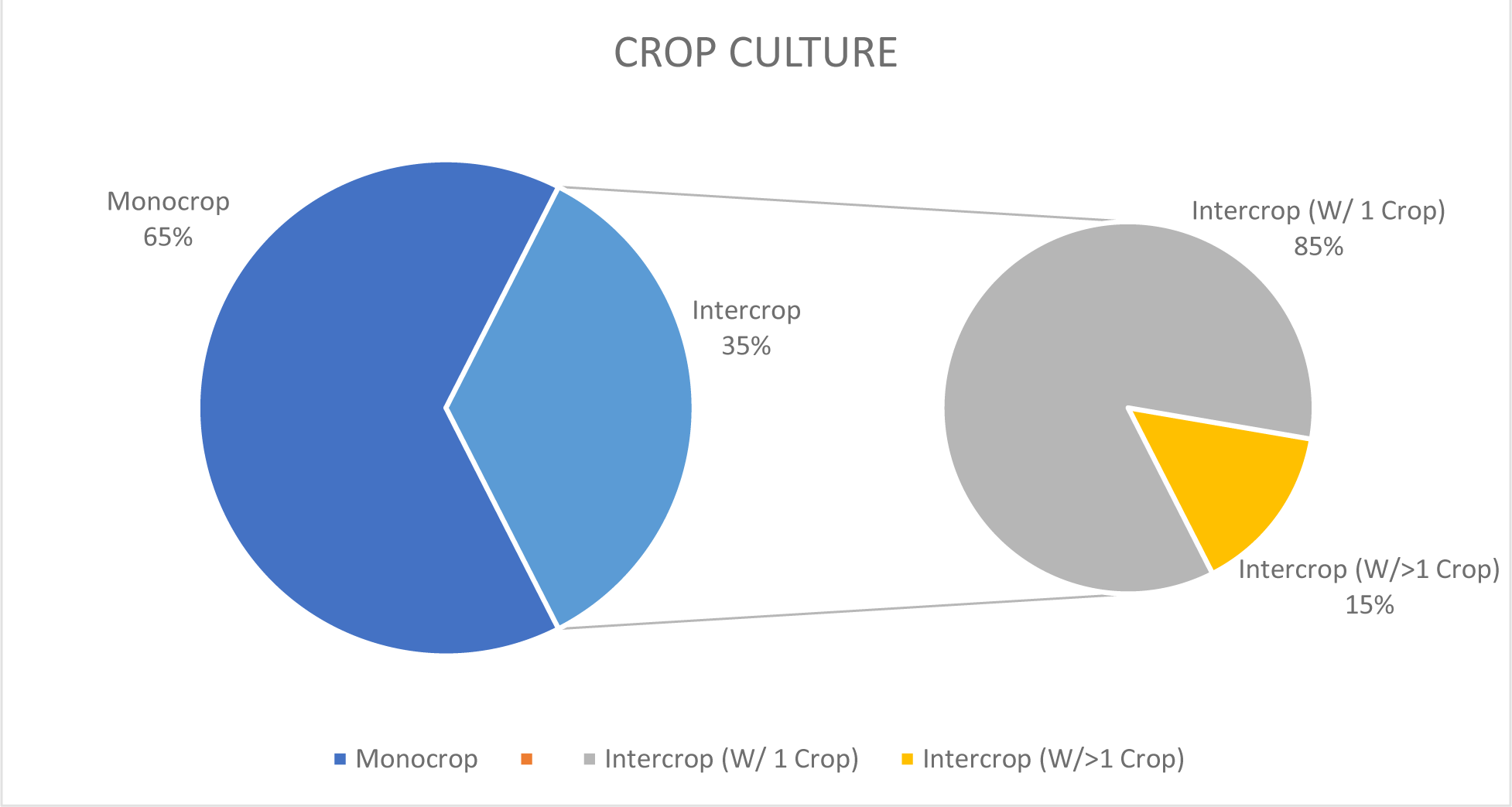
Composition of fields surveyed in Busia, Kenya between August 12^th^ and November 21^st^, 2019.

The distinction between intercropped with one other crop versus intercropped with two or more other crops was made due to the low population of intercropped fields with two or more other crops (15%) out of intercrop field composition. The combinations of intercropped fields with two or more other crops is broken down further in Table 3.

**Figure 4:**
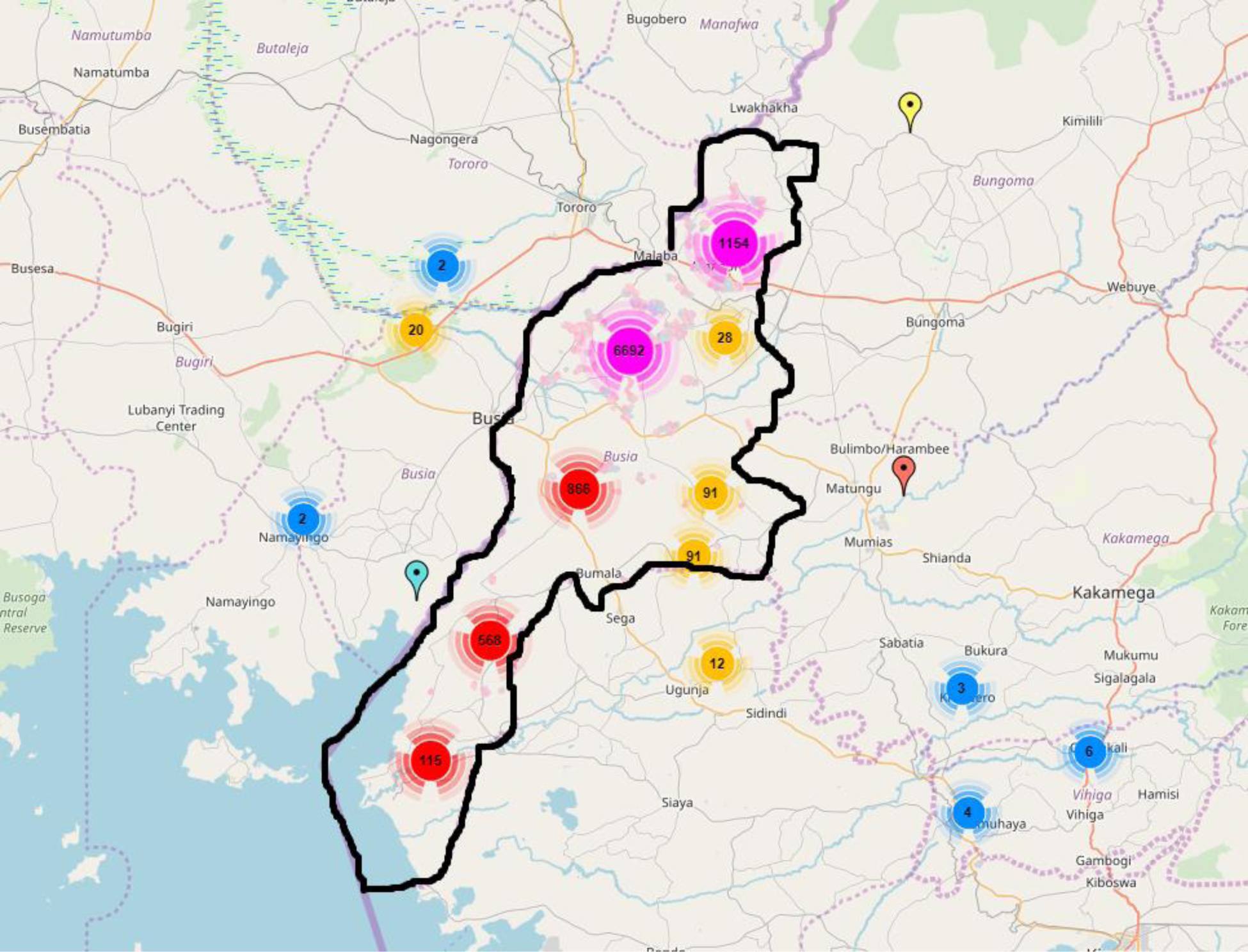
Distribution of the field collections and disease detections across Busia county as of October 14^th^, 2019.

The analysis of monocrop fields can be found in Table 1. The ‘Percentage of Fields’ column describes the proportion of all monocrop fields that are made up of each specific crop. Table 1 shows the analysis of intercrop fields with one other crop. The ‘+’ dictates the crop listed plus any other crop. The ‘Percentage of Fields’ column under intercropped fields describes the percentage of the crop plus any other crop out of all of the fields intercropped with one other crop.

**Figure 5:**
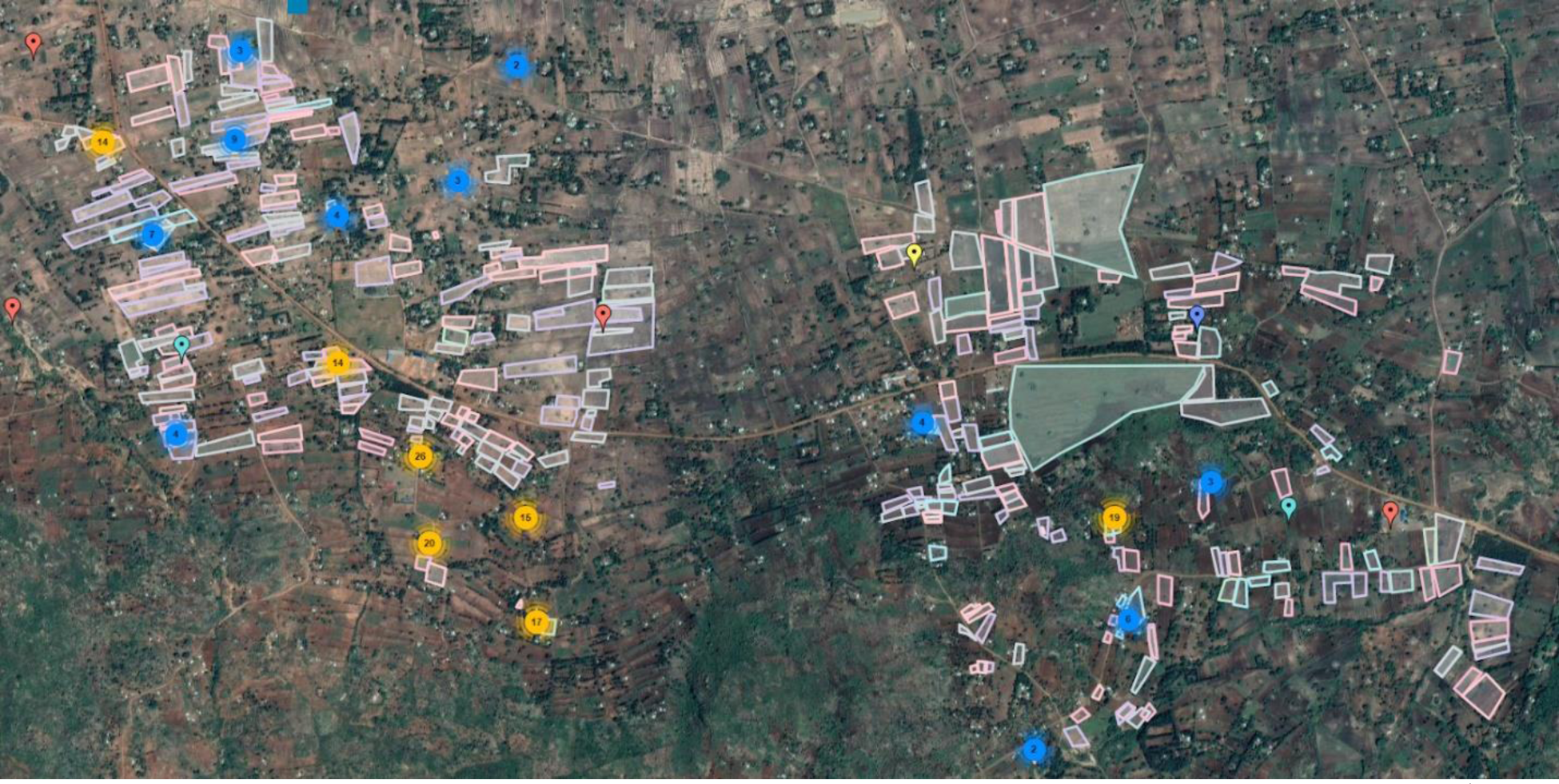
Satellite image from PlantVillage website showing field collections and disease detection data as of October 14^th^, 2019

**Table 1:**
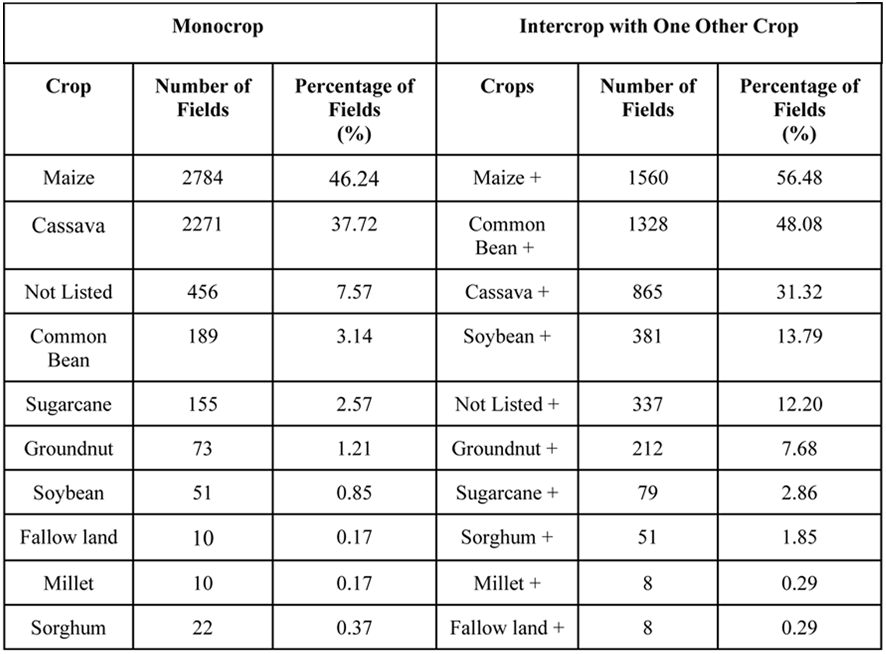
Detailed composition of the fields surveyed between August 12 and October 08, 2019 in Busia, Kenya

**Table 2:**
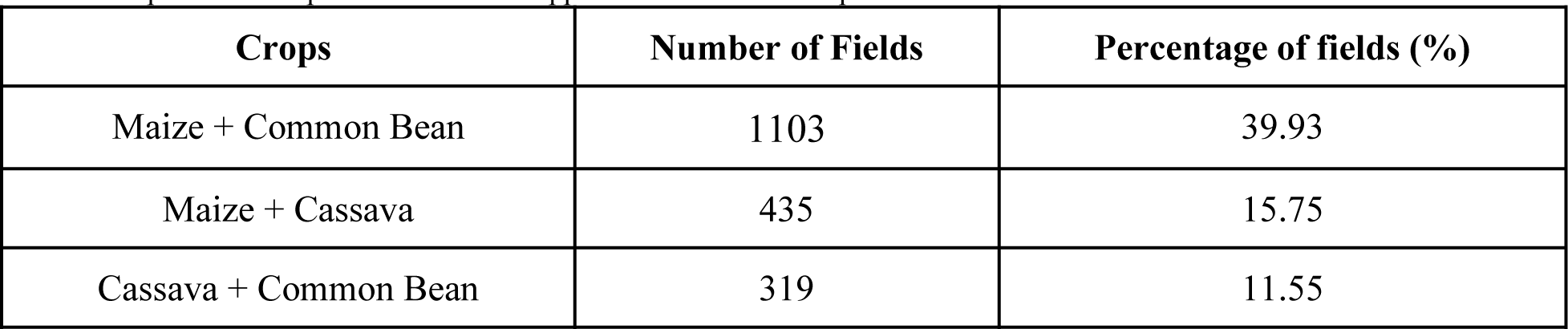
Top 3 field compositions of intercropped fields with one crop.

**Table 3:** Number of fields intercropped with two or more other crops.

| **Crops** | **Number of Fields** | **Percentage of fields (%)** |
| --- | --- | --- |
| With 2 other crops | 423 | 90.00 |
| With 3 other crops | 44 | 9.36 |
| With 4 other crops | 3 | 0.64 |

### Dynamic Interface

The following images were captured from the PlantVillage website on October 14^th^, 2019. These images include fields surveyed represented as various colored polygons. The colors represent the main crop in the field. The markers represent disease diagnosis from the app PlantVillage *Nuru* and the WaPOR evapotranspiration data is represented as green, amber or red circles. A green circle indicates the crop is not stressed which is determined by reference evapotranspiration equaling actual evapotranspiration. An amber circle means your crop is minimally stressed, which occurs when your reference evapotranspiration is minimally less than your evapotranspiration. A red circle means your crop is stressed and the reference evapotranspiration is below the actual evapotranspiration. Further explanation are provided on the site and the color distinctions may change following input from agronomists studying crop responses to water stress.

**Figure 6:**
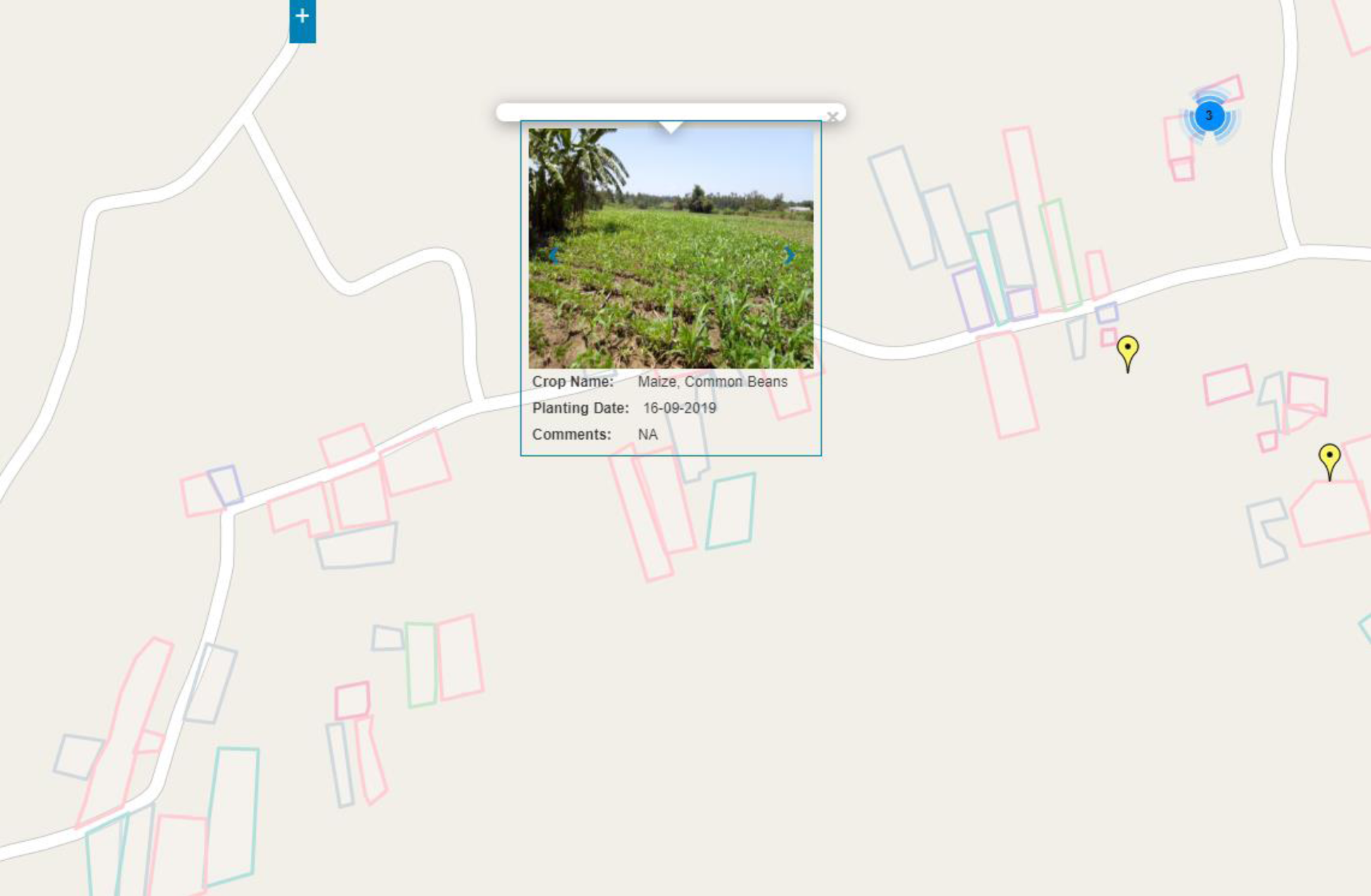
Image from PlantVillage website showing field collection data and disease detections as of October 14^th^, 2019.

### Crop Stress via WaPOR

The WaPOR platform can be used to determine crop water stress. This can be determined from the value of the Actual Evapotranspiration (Allen et al-998). As this is an evolving product that is intended to be built as a climate change adaptation tool through collective action of the scientific community we are not presenting results here.

**Figure 7:**
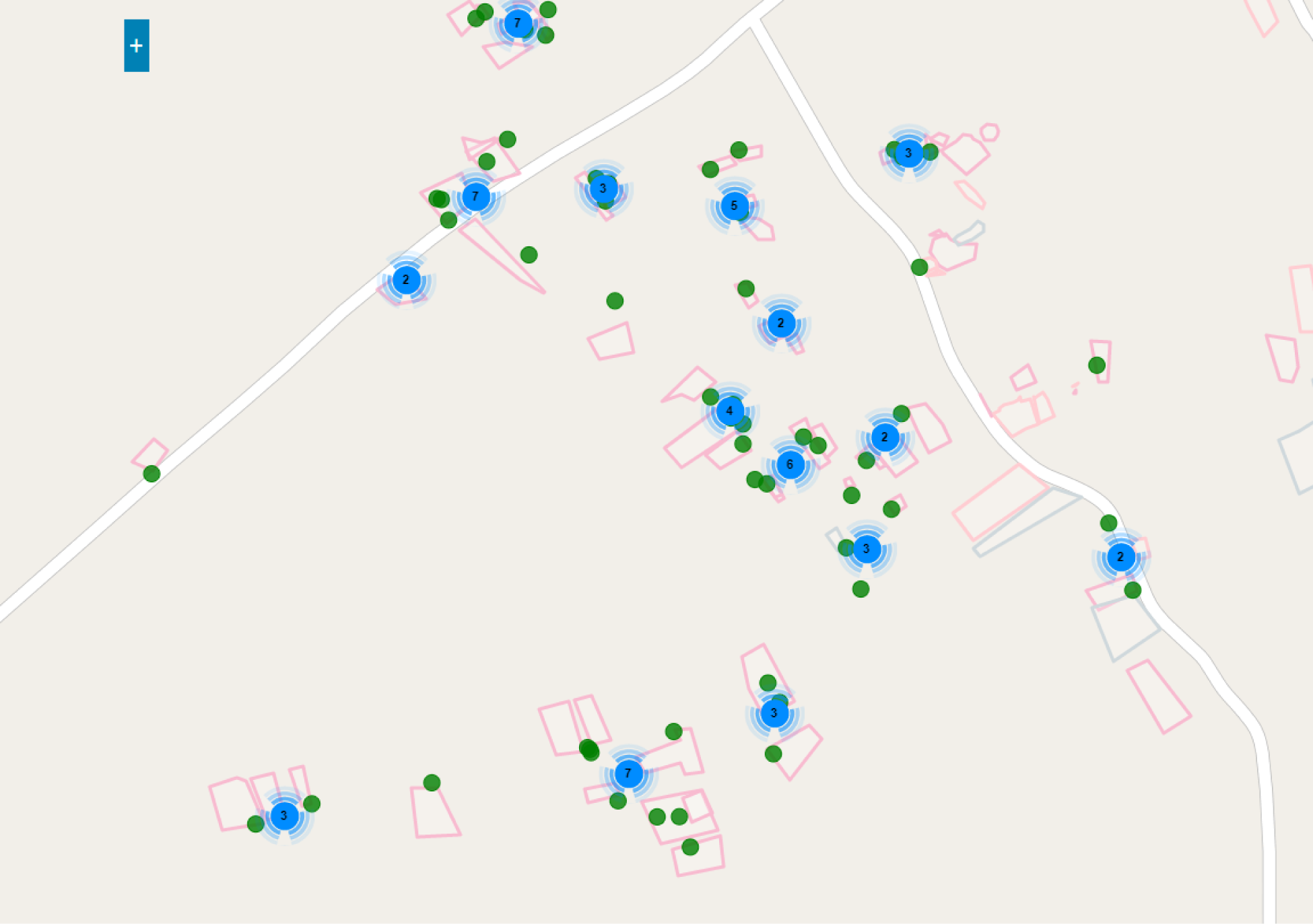
Image from PlantVillage website showing WaPOR crop stress for field collections as of October 14^th^, 2019.

## Discussion and Conclusion

### Data Collection Rates and Limitations

The average collections rate for the Kenyan survey team is 128 fields per day. The team captured field composition data for 9,263 fields during August 13^th^ and November 21^st^, 2019. If the team were to maintain this rate, they could survey approximately 36,000 fields in one year. The frequency at which this data can be collected compared to the cost of collection (approximately $1.41 USD per field) is extremely favorable. By providing motivated University students and recent graduates with the means to earn a steady income, help their community, and support a movement for climate change adaptation, the status of Kenya’s agriculture can be dramatically enhanced a low cost.

A limitation we currently face is not knowing the number of fields needed to sufficiently represent a region. Each region can have differing spectral profiles for the same crop on remotely sensed images due to factors like weather (Foerster-2012). If the number of fields required to represent a region can be determined, the field data collection model can be optimized across all counties in Kenya and ultimately scaled across Africa.

### Field Data Collection and Challenges

The field compositions for season one was mainly monocropped maize (39%) and monocropped cassava (14%). We found the main reason farmers planted maize instead of cassava in season one was due to the diseases that affect cassava and its cuttings. Another reason for planting maize is that cassava seeds are more expensive than maize seeds and the farmer is not guaranteed to get the variety they think they are purchasing. We found that a large number of maize fields in season one were replanted two to three separate times due to delay in the onset of seasonal rains (GEOGLAM-2019) which in March/April and was associated with the twin Cylones in Mozambique (Idai and Kenneth) which removed moisture from the Horn of Africa (GEOGLAM-2019).

The survey team found that maize (46.24%) and cassava (37.72%) were the majority of monocrop fields in season two. However, there was an increase in fields intercropped with common bean from 21.8% in season one to 48.08% in season two. This is due to the shorter rains during the second season, which reduces the risk of fungal diseases in beans according to the farmers.

The Kenyan survey team faced and continues to face several challenges while surveying fields. One such challenge is the variability in data connection while in the field. If the data connection is poor, the satellite image will not load in the ODK app making it impracticable to determine the boundaries of the field from the satellite imagery. When this is the case, the Kenyan survey team must walk the boundaries of the field for every field. This is physically taxing and takes time so when this approach is required, the total number of fields surveyed for the day decreases but the quality of the mapping increases as more field boundary data points are collected with this method.

Another challenge the survey team faced and overcame was reaching farmers and fields outside of the SHA network. The team exhausted the farmers and fields within the SHA network in Busia after just a few weeks. They overcame this challenge by meeting with community groups that meet regularly to engage new farmers and reach previously existing networks that are not part of SHA. The added value of this was an ability to share more details on pests and diseases as well as adaptive approaches to coping with changes in weather associated with climate change.

The fields collected are updated daily^3^ for everyone to see and download. We understand the importance of this data and how applicable it is to various fields and therefore offer it on the PlantVillage website for free to encourage others to download and interpret this data using their expertise. On the site, the user can see the fields surveyed and collected information from the Kenyan survey team along with NURU diagnoses and WaPOR data for the points. The combination of WaPOR data with field composition data can tell the user in real-time the status of each field. This is critically important in determining the advice given to the farmers.

### Crop Stress as a Function of Field Composition Data and WaPOR

Field composition data are used in combination with WaPOR evapotranspiration data to create a crop stress map across Busia county. This is a work in progress and is being released as a alpha stage product to encourage the global community of agronomists to work collaboratively on developing machine learning approaches to provide near real-time advice to smallholder farmers on climate stress.

The seasonal rains that were expected in March did not arrive until early April, which was due in large part to the twin Cyclones in Mozambique (GEOGLAM-2019). This caused a large shift in either later planting dates or low-germination rates. The effect of climate change is likely to result in a greater shift in planting dates. This implies that a dynamic tool that provides a more accurate index of when to plant would offer significant value. As part of our efforts at PlantVillage, we are developing a climate-smart Artificial Intelligence (A.I.) Assistant which integrates the ground-truthed data presented with evapotranspiration data from WaPOR to help farmers adapt to the significant challenges ahead.

### Conclusion

The capability of young motivated Moi University graduates created a high-throughput system for accurate data collection in agriculture. This pilot study demonstrates it is economically viable to expand and sustain youth-led teams to improve the agricultural system in Kenya and other countries in Africa.

## Data Availability Statement

All datasets described in this study are available online at www.plantvillage.psu.edu/AfricaAI and https://wapor.apps.fao.org/home/WAPOR_2/1.

## Author Contributions

AK, DH and JC conceived the project. AK designed the initial field survey protocol. AK, PM, JC, SA, BP, JM, GN, KN, LP, MJ, JM, MT, WO collected the data. AK analyzed the data collected. DM analyzed WaPOR data. JC hired and managed Kenyan survey team. AK, DH, JC contributed to writing. AK, PM, JC, DH contributed to editing.

## Acknowledgements

We are very grateful to the many farmers who welcomed us onto their land and allowed us to collect this data for the common good. We are grateful to Fabio Lana for technical input and Hidden Brains for development work on PlantVillage. We thank Self Help Africa for local support and Inbal Becker-Reshef for inviting this contribution.

## Supplementary Material

Table S1, Table S2, Table S3

## Footnotes

1 *Dove - Satellite Missions - EoPortal Directory*, earth.esa.int/web/eoportal/satellite-missions/d/dove.

2 “Leader in Satellite Imagery.” *DigitalGlobe*, www.digitalglobe.com/company/about-us.

3 *AfricaAI* -*PlantVillage*. www.plantvillage.psu.edu/AfricaAI

4 *WaPOR – FAO*, https://wapor.apps.fao.org/home/WAPOR_2/1

5 *Open Data Kit – Build –* build.opendatakit.org

